# Fecal filtrate transfer protects against necrotizing enterocolitis in preterm pigs

**DOI:** 10.1101/2020.05.25.114751

**Authors:** Anders Brunse, Ling Deng, Xiaoyu Pan, Yan Hui, Witold Kot, Duc Ninh Nguyen, Jan Bojsen-Møller Secher, Dennis Sandris Nielsen, Thomas Thymann

**Affiliations:** Department of Veterinary and Animal Sciences, Faculty of Health and Medical Sciences, University of Copenhagen, Denmark; Department of Food Science, Faculty of Science, University of Copenhagen, Denmark; Department of Plant and Environmental Sciences, University of Copenhagen, Denmark; Department of Veterinary Clinical Sciences, Faculty of Health and Medical Sciences, University of Copenhagen, Denmark

## Abstract

**Background and aims:** Necrotizing enterocolitis (NEC) is an acute and life-threatening gastrointestinal disorder afflicting preterm infants, which is currently unpreventable. Fecal microbiota transplantation (FMT) is a promising preventative therapy, but potential side effects raise concern. Removal of bacteria from donor fecal water may reduce side effects while maintaining wanted effects. We aimed to assess preclinical efficacy and safety of bacteria-free fecal filtrate transfer (FFT).

**Methods:** Using fecal material from healthy suckling piglets, we administered rectal FMT or cognate FFT by either rectal or oro-gastric administration to formula-fed preterm, cesarean piglets, and compared gut pathology and related safety parameters with saline controls. We then analyzed mucosa and luminal bacterial and viral composition using 16S rRNA gene amplicon and metavirome sequencing, respectively. Finally, we used isolated ileal mucosa, coupled with RNA-Seq, to gauge the host response to the different treatments.

**Results:** Oro-gastric FFT eliminated NEC, which was confirmed by microscopy, whereas FMT did not perform better than control. Moreover, FFT but not FMT reduced intestinal permeability, whereas FMT animals had reduced body weight increase and intestinal growth. Oro-gastric FFT increased viral diversity and reduced Proteobacteria abundance in ileal mucosa relative to control. Global gene expression of host mucosa responded to FMT but not FFT with increased and decreased bacterial and viral defense mechanisms, respectively.

**Conclusions:** As preterm infants are extremely vulnerable, rational therapies need incontestable safety profiles. Here we show in a clinically relevant animal model that FFT, as opposed to FMT, efficiently prevents NEC without any recognizable side effects. If translatable to preterm infants, this could lead to a change of practice and in turn a reduction in NEC burden.

## INTRODUCTION

Gut colonization after birth is essential for development of the host immune system. Yet, it has become increasingly clear that microbial perturbation resulting in deviation from the normal gut microbiota developmental trajectory is a risk factor and possibly a contributing factor for a range of neonatal diseases. Gut dysbiosis is of particular concern among preterm infants due to their impaired microbial host defenses and high susceptibility to life-threatening infections^1^. Moreover, preterm infants are often born by cesarean section^2^, receive extensive amounts of antibiotics^3^, and endure invasive procedures, similarly contributing to gut microbiota perturbation and infection risk^4^.

Necrotizing enterocolitis (NEC), a lethal inflammatory and necrotic bowel disease mainly affecting very preterm infants, is a prominent example of a disease that is tightly coupled with gut dysbiosis^5^. Under experimental conditions, germ-free rearing and enteral antibiotics treatment provide complete protection from NEC, demonstrating the necessity for gut microbiota presence and bulk in NEC etiology^6–8^. Moreover, preterm infants generally respond well to probiotics treatment, which appears to offer some protection against NEC^9^. In preterm infant cohorts, temporal and geographical differences^10^ complicate the expression of a generalized NEC microbiota, although increased Proteobacteria and reduced Bacteroidetes abundance are unifying features of a prediagnostic NEC microbiota across neonatal units^11^. At single sites, the microbiota preceding NEC diagnosis is usually characterized by reduced bacterial diversity, lack of obligate anaerobes and overabundance of single facultative anaerobes often belonging to the family of Enterobacteriaceae^12–14^.

In the search for better bacterial therapies, we recently showed a proof of principle for fecal microbiota transplantation (FMT) against NEC, using cesarean-delivered preterm pigs as recipients and breastfed, term pigs as donors^15^. Whereas the beneficial effect of FMT on gut pathology was unequivocal, and the engraftment of perceived beneficial donor bacteria into the recipients significant, the safety profile raised some concern, chiefly by increasing sepsis incidence and mortality, when FMT was administered orally. Besides bacteria, the fecal matrix consists of archaea, eukarya, viruses, microbial secretome and metabolome, any of which may be attributable to the benefits and adversities of FMT. Accordingly, any means to reduce the complexity of the donor fecal matrix, while maintaining its therapeutic effects is an advancement towards developing a clinically feasible therapy against NEC.

Bacteriophages, viruses infecting bacteria in a host-specific fashion, are omnipresent across all bacteria-containing ecosystems including the mammalian gut. During early gut colonization bacteriophages and bacteria dynamically interact and influence each other’s composition and abundance^16,17^. Interestingly, a small case series of *Clostridioides difficile* infection patients receiving bacteria-free donor fecal filtrate transfer (FFT) reported cure rates equivalent to regular FMT treatment and attributed the effect to donor bacteriophages^18^. Accordingly, donor bacteriophages might mediate the beneficial effect of FMT on NEC. Further, it has recently been shown that FFT from lean mice to mice on a high fat diet reduces weight gain and protects against metabolic syndrome development in the recipients^19^, again indicating that FFT is able to modulate the recipient phenotype.

We hypothesized that sterile fecal filtrate transfer (FFT) would be safe and non-inferior to FMT for NEC prevention. Using cesarean-delivered preterm pigs as models for very preterm infants, we compared the clinical and gut microbiological effects of FFT by different routes of administration with FMT and control. To our surprise, we found that orally administered FFT was superior to FMT by offering almost complete protection against NEC. Furthermore, FMT-associated adverse effects did not occur in relation with FFT treatment. Concurrently, FFT changed the gut viral and bacterial composition in an administration route-dependent manner. In summary, we discovered FFT as a safe and effective means to reduce NEC incidence.

## MATERIALS AND METHODS

### Animal experimental procedures

The Danish Animal Experiments Inspectorate approved all experimental procedures (2014-15-0201-00418). Seventy-three conventional crossbred piglets (Landrace x Large white x Duroc) from three healthy sows were delivered by caesarean section at 90% gestation. Birth, resuscitation and housing conditions are described in details elsewhere^15^. Animals were fitted with oro-gastric feeding tube and an intra-arterial catheter through the transected umbilical cord, and received an infusion of 16 ml/kg maternal plasma to compensate for the lack of transplacental immunoglobulin transfer. We then stratified the animals by sex and birth weight, and randomly allocated them to four groups receiving rectal fecal microbiota transplantation (FMT), rectal fecal filtrate transfer (FFTr), oro-gastric fecal filtrate transfer (FFTo) or oro-gastric and rectal control saline (CON). All animals received increasing volumes of infant formula by tube feeding (24 – 96 ml/kg/d, composition in Supplementary table S1) with decreasing parenteral nutrition supplement (96 – 48 ml/kg/d, Kabiven, Fresenius Kabi, Copenhagen, Denmark).

### Fecal microbiota transplantation and fecal filtrate transfer

The donor fecal material used in this experiment originated from the same batch that previously showed a beneficial effect against NEC^15^. Briefly, colon luminal content was collected from five 10-day-old piglets and then pooled, gently homogenized and frozen in 10% sterile glycerol. For the FMT solution, thawed fecal material was diluted in sterile saline to 0.05 g/ml and filtered through a 70-μm cell strainer. The FFT solution was prepared in advance as previously described^20^. Briefly, thawed fecal material was diluted to the same concentration as above, homogenized, centrifuged at 5000x g for 30 min at 4 °C, and supernatant filtered through a 0.45 μm PES filter (Minisart^®^ High Flow Syringe Filter, Sartorius, Göttingen, Germany). Twice daily on days 1 and 2, animals received 0.5 ml treatment solution. Rectal administration (CON, FMT, FFTr) was performed with a soft rubber probe placed 3-5 cm into the rectum, whereas oro-gastric solutions (CON, FFTo) were administered in the feeding tube followed by flushing with 1 ml of sterile water.

### Clinical monitoring and euthanasia

Animals were monitored by experienced personnel, and daily weights and stool patterns recorded. Animals presenting with clinical signs of NEC or systemic illness throughout the experiment were immediately euthanized. On day 5, animals were deeply anesthetized and euthanized with a lethal cardiac injection of barbiturate. Three hours prior, animals received 15 ml/kg of a 5/5% w/v lactulose and mannitol solution in the feeding tube to measure intestinal permeability. After abdominal incision, urine was collected by cystocentesis for lactulose and mannitol measurement as previously described^21^. Abdominal organs were excised and weighed, and gross pathological changes to the stomach, small intestine, and colon were assessed in accordance with an established six-grade NEC scoring system^15^. The highest grade assigned expressed the disease severity, and NEC diagnosis was defined as pathology grade 4 (extensive hemorrhage). Luminal content was collected from ascending colon for gut microbiota analysis. Three biopsies were collected along the small intestine for lactase activity measurement^21^. Biopsies of distal ileum and transverse colon were fixed in paraformaldehyde and later embedded in paraffin, sectioned and stained with hematoxylin and eosin for histopathological evaluation. Tissue sections were blindly evaluated based on a six-grade scoring system (Supplementary figure S1), where microscopic NEC was defined as histopathology grade 4 or above. Finally, a 10 cm section of distal ileum was inverted, washed in sterile saline and blotted to remove residual fluid. Mucosa tissue was then scraped off using sterile object glass and cryopreserved for gut microbiota analysis and host transcriptomic analysis.

### Gut microbiota composition

The bacterial compositions of distal ileal mucosa and gut luminal content were determined by 16S rRNA gene (V3-region) amplicon sequencing on a NextSeq using PE150 (Illumina, San Diego, CA, USA). Total DNA was extracted using Bead-Beat Micro AX Gravity Kit (A&A Biotechnology, Gdynia, Poland) according to the manufacturer’s instructions. Library preparation followed a previously published protocol^22^. The raw sequencing reads were merged and trimmed. Chimeras were removed and zero-radius Operational Taxonomic Units (zOTUs) constructed using UNOISE algorithm implemented in Vsearch^23–25^. The Greengenes (version 13.8) database was used as reference for annotation. Qiime2 was used to process the forward analysis. Rare zOTUs with frequency below 0.1% of the minimal sample depth were filtered and removed, and based on rarefaction curve the zOTU table was rarified to adequate sample depth (9000 counts) for alpha and beta diversity calculations. Principal coordinate analysis (PCoA) was conducted on unweighted UniFrac dissimilarity metrics, a PERMANOVA test with FDR correction was performed to detect pairwise group differences. Specific taxa comparison among groups was analyzed by ANCOM with default Qiime2 settings used to test statistically significant differences.

The viral content from the same gut mucosa and luminal samples were purified, DNA extracted and library constructed, sequenced and analyzed as previously described^19,20^. The average sequencing depth for the viral metagenome was 6,158,777 reads/sample (min. 72,077 reads and max. 13,788,165 reads). For each sample, reads were treated with Trimmomatic^26^ and Usearch^27^ and subjected to within-sample de novo assembly with MetaSpades v.3.5.0^28,29^. Viral contigs were identified with Kraken2^30^, VirFinder^31^, PHASTER^32^, and virus orthologous proteins (www.vogdb.org). Contaminations of non-viral contigs were removed. The remaining contigs constituted the vOTU (viral-operational taxonomic unit) table. Analysis of viral community α- and β-diversity were performed using packages Phyloseq v1.30.0 and Vegan 2.5-6 in R. For α-diversity analysis, Shannon index was calculated. The contig table of viral data was normalized by counts per length of contig (kb) per million reads mapped (TPM). Bray-Curtis distance metrics were calculated for β-diversity analysis. Unconstrained ordination was performed using principal coordinate analysis (PCoA). Constrained ordination was done using distance-based redundancy analysis (dbRDA, ‘capscale()’ function in Vegan). Significant predictor variables were identified by PERMANOVA tests. Results were visualized in graphical forms using a set of custom R scripts and functions in the packages of Phyloseq, Vegan and ggplot2.

### RNA seq

Global trancriptomic patterns were investigated in RNA extracts from terminal ileal mucosa by RNA seq. Total RNA was isolated with RNeasy Micro Kit (Qiagen), and 1.5 μg RNA per sample was used for library construction. Sequencing libraries were constructed using NEBNext Ultra RNA library Prep Kit for Illumina (New England Biolabs, Ipswich, MA, USA) and sequenced on the Illumina HiSeq 4000 platform with paired-end 150-bp reads production. Quality and adapter trimming of raw reads was performed using TrimGalore (Babraham Binoinformatics). Alignment against the porcine reference genome Sscrofa11.1 was performed with the RNA-seq aligner Tophat^33^. Gene counts were obtained with HTSeq-count^34^, using gene annotations from the Ensembl genome assembly (v91). Statistical analysis of differential gene expression between groups was performed by DESeq2^35^ in R statistical software using an FDR adjusted cutoff of p<0.10 and log2 fold change > 1. Analysis of gene-enrichment and functional annotation was performed with Cytoscape (version 3.7.1) using an FDR adjusted cutoff of p<0.10.

### Systemic immune cell characterization

Complete blood cell counts and basic T cell phenotyping was performed in blood samples collected on days 3 and 5, as previously described^36^.

### Statistics

NEC scores and urinary lactulose-mannitol ratio were analyzed by Kruskal-Wallis tests. NEC, rectal bleeding and growth failure incidences were analyzed by Fisher’s exact tests. Lactase activity was analyzed by two-way ANOVA, and remaining continuous data were analyzed by one-way ANOVA. Probability levels below 0.05 were considered significant.

## RESULTS

### Initial clinical course

Among the 75 cesarean-delivered preterm piglets, nine were removed before randomization (failed resuscitation, stillbirth, intrauterine growth-retardation and infection), whereas the remaining 66 animals were group allocated. An additional seven animals were euthanized preschedule of non-gastrointestinal causes (respiratory failure, iatrogenic complications) and removed from the experiment. Two animals were euthanized preschedule with clinical NEC signs (1 CON, 1 FFTr), whereas the remaining 57 animals survived until day 5. During the course of the experiment, we observed rectal bleedings in 31% (5/16) of CON and 19% (3/16) of FMT animals relative to 0% (0/13) in both FFT groups (p < 0.05 vs. CON).

### Gut pathological evaluation

We performed gross examination of the gastrointestinal tract supported by histopathological assessment of ileum and colon to evaluate the severity and extent of NEC-like lesions in response to the different treatments. The NEC-like pathological phenotype in formula fed preterm pigs consisted of extensive hemorrhage with or without patchy necrosis of the mucosa and pneumatosis intestinalis, mostly affecting ascending and transverse colon, and to a lesser extent ileum and stomach (Fig. 1A). On the microscopic level, the pathological observations included subtle changes to the mucosal architecture, progressing from epithelial sloughing and hemorrhage to complete destruction of mucosal integrity among the most severe cases (Fig. 1B).

**Figure 1.**
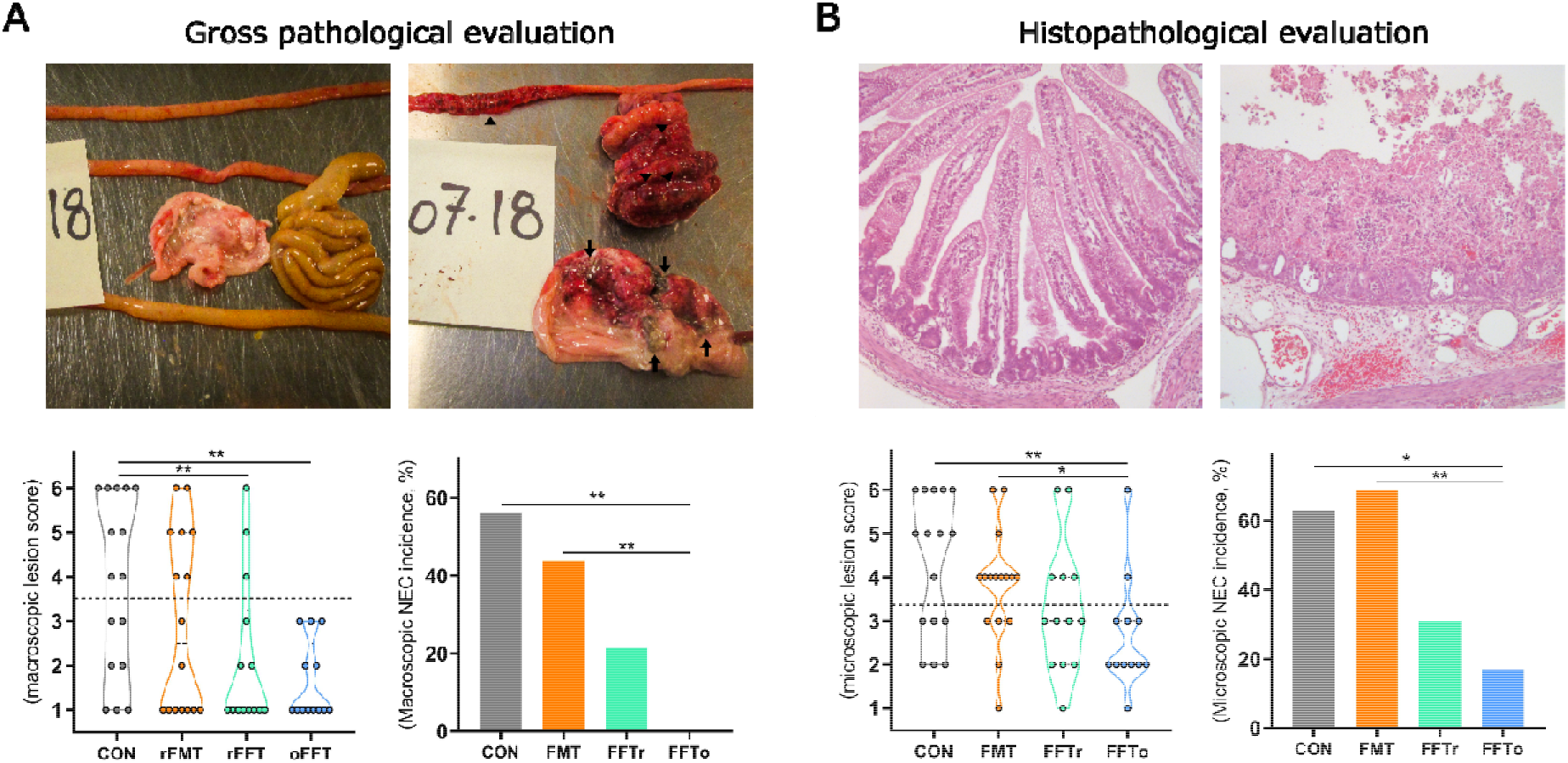
Gut pathological evaluation following FMT and FFT treatment. A. Representative necropsy photographs of pig stomach, small intestine and colon with minimal (upper left) or severe pathological changes (upper right). Arrows point to necrotic patches in the mucosa, and arrowheads highlight macroscopic pneumatosis intestinalis. Pathological severity (lower left) and macroscopic NEC incidence (lower right) are showed. B. Representative micrographs of hematoxylin & eosin stained intact (upper left) and severely disrupted small intestine (upper right) captured at 10x magnification. Histopathological severity (lower left) and microscopic NEC incidence (lower right) are showed. *, ** denote statistical probability levels of 0.05 and 0.01, respectively.

In this setting, oro-gastric FFT administration markedly reduced NEC severity (p < 0.01, Fig. 1A) and consequently reduced NEC incidence to 0% (p < 0.01 vs. CON). Rectally administered FFT also reduced NEC severity relative to CON (p < 0.01), but due to three NEC diagnoses in this group, the reduction in NEC incidence did not reach statistical significance (p=0.07 vs. CON). However, rectally administered FMT, which we previously found to be clearly NEC protective^15^, failed to reduce NEC severity and incidence relative to CON. Of note, no bleeding occurred after rectal fluid administration, and no rectal lesions were observed at necropsy in rectally administered animals. The microscopic evaluation supported the macroscopic effects of oro-gastric FFT, which reduced the histopathological NEC severity and incidence relative to CON as well as FMT (all p > 0.05, Fig. 1B), whereas no significant effects were found for rectally administered FFT on microscopic level. Notably, animals receiving FMT had a characteristic pattern of epithelial sloughing in distal ileum, which was rarely observed in any other group.

### Safety assessment

As an integral part of preclinical evaluation and due to concern about the safety of FMT and FFT to preterm neonates, we next investigated a series of safety parameters. Initially, we found that several animals receiving FMT had a negative body growth rate (Fig. 2A). Hence, the FMT group had a significantly higher proportion of animals with a negative body growth rate compared with CON and FFTo (both p < 0.05, Fig. 2B). Furthermore, the relative weight of the small intestine but not colon was lower in FMT animals relative to FFTo (p < 0.05, Fig. 2C). Additionally, the urinary lactulose-mannitol ratio, a dynamic *in vivo* marker of small intestinal permeability, was robustly decreased by FFTr (p < 0.05 vs. CON, Fig. 2D), whereas the FMT group had a significantly higher permeability than both FFT groups (both p < 0.05). Finally, the lactase enzymatic activity across the small intestine, a measure of mucosal integrity, was significantly decreased in the FMT group relative to FFTr (Fig. 2E).

**Figure 2.**
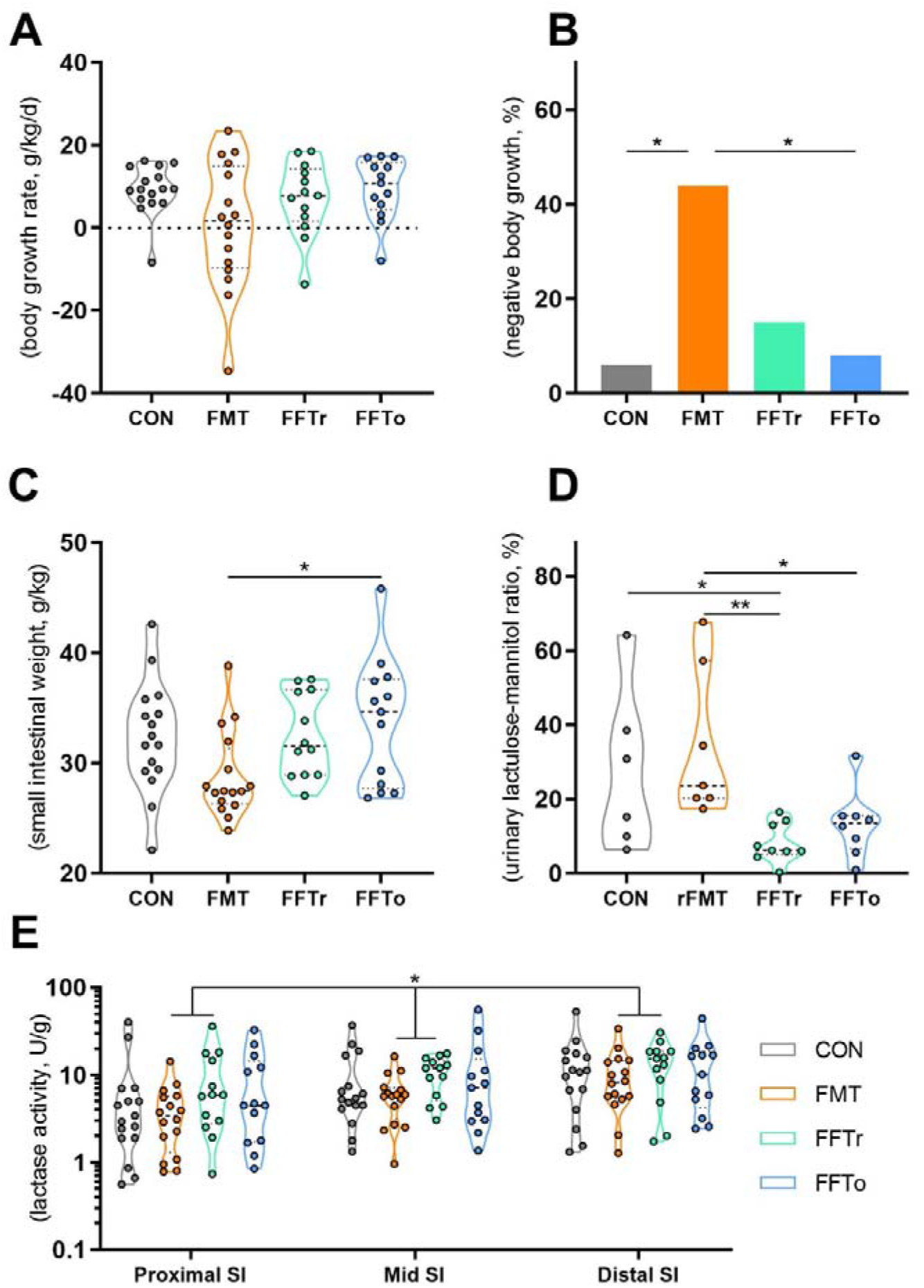
Safety assessment data following FMT and FFT treatment. A. Relative body growth rate from birth to day 5. B. Proportion of animals with negative body growth rate from birth to day 5. C. Relative small intestinal weight. D. Small intestinal permeability as assessed by the ratio of lactulose and mannitol recovered in urine after a standardized oro-gastric administration. E. Brush-border lactase enzyme activity in three segments of small intestine. SI, small intestine; *, ** denote statistical probability levels of 0.05 and 0.01, respectively.

### Gut mucosal and luminal microbiota composition

We next investigated how mucosa-associated and gut luminal bacterial and viral communities changed in response to the different treatments. Across all animals, the composition of mucosa-associated bacteria differed from luminal bacteria (Unweighted UniFrac, R2 = 0.11, p < 0.001, Supplementary figure S2). Intervention effects were seen in bacterial composition of both mucosa (R2 = 0.17, p < 0.001, Fig. 3A) and luminal compartments (R2 = 0.27, p < 0.001). The greatest effect was observed in the FMT group, but both FFT groups were also significantly different from CON in both mucosa and gut lumen. However, FFT route of administration (FFTr vs FFTo) did not affect the bacterial composition (Supplementary table S2). The Shannon diversity index of mucosa-associated bacteria was similar among groups, but interestingly a positive correlation between Shannon diversity index and small intestinal NEC severity was observed (R^2^ = 0.41, p < 0.001, Supplementary figure S3). The luminal bacterial Shannon diversity index increased only in response to FMT treatment (Fig. 3B).

**Figure 3.**
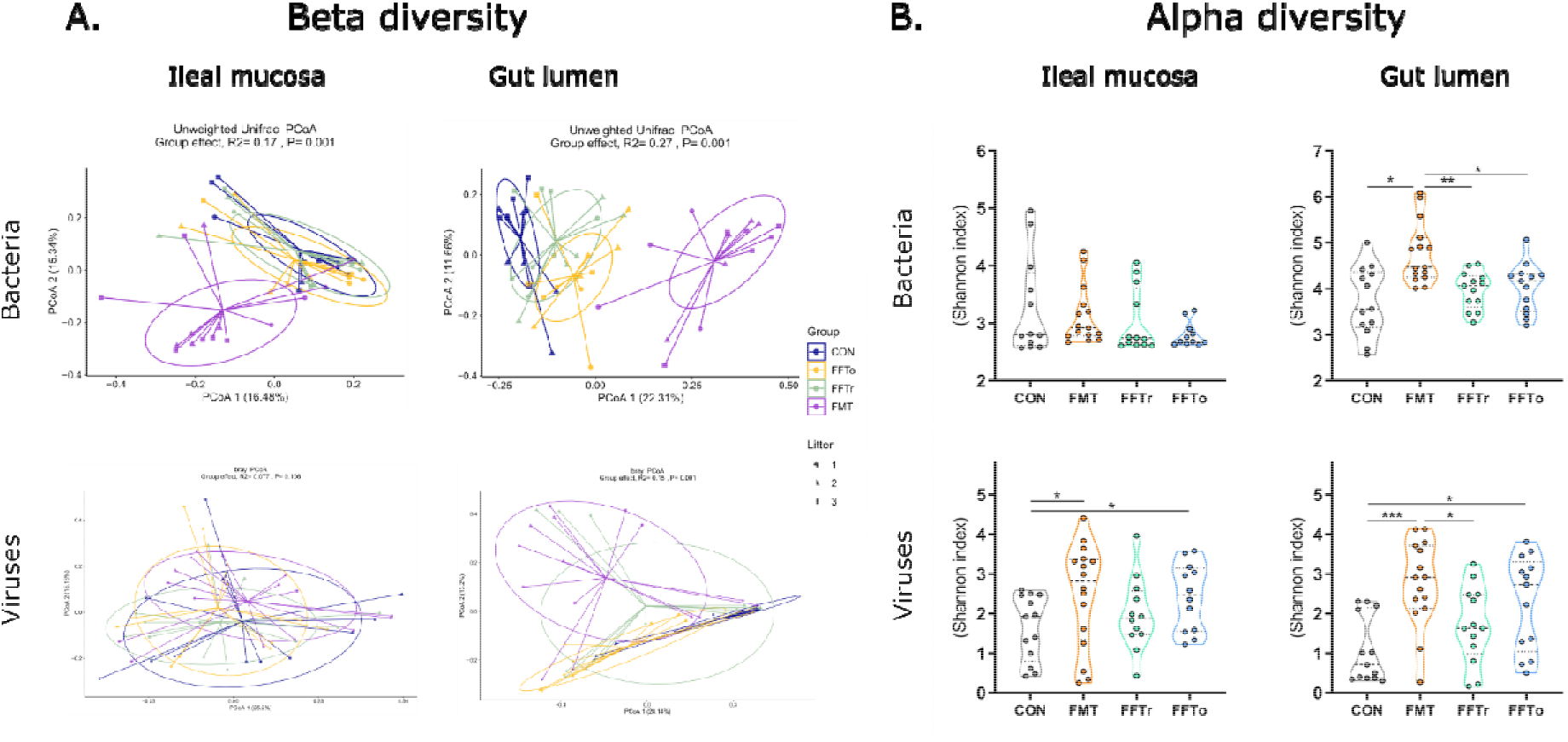
Gut mucosa and luminal bacterial and viral composition following FMT and FFT treatment. A. Principal component analysis plots visualizing bacterial (upper) or viral (lower) beta diversity among groups in the mucosal (left) or luminal (right) niche. Bacterial beta diversity is based on unweighted UniFrac mettrics, whereas viral beta diversity is based Bray-Curtis dissimilarity metrics. B. Shannon index as a measure of bacterial (upper) or viral (lower) alpha diversity among groups in the mucosal (left) or luminal (right) niche. *, **, *** denote statistical probability levels of 0.05, 0.01 and 0.001, respectively.

As previously observed, the bacterial microbiota was dominated by Enterococci (>70% abundance across groups and anatomical sites)^15^. Remarkably, two genera accounted for the majority of differences in the residual composition. Whereas the donor feces contained high levels of both *Streptococcus* and *Lactobacillus* and CON animals harbored neither, FFT treatment substantially increased *Streptococcus* abundance but not *Lactobacillus*, while the opposite was true for FMT (Supplementary figure S4). Interestingly, while the abundance of Proteobacteria was similar among groups in the gut lumen, very few Proteobacteria were detected from mucosa of FFT treated animals, whereas approximately 10% (relative abundance) of CON mucosa consisted of Proteobacteria (e.g. Enterobacteriaceae, *Klebsiella)* and FMT approximately 5%.

Treatment effects on viral composition were observed in the gut lumen (Bray-Curtis dissimilarity, R2 = 0.15, p = 0.001, Figure 3A), but not the mucosa compartment (R2 = 0.077, p = 0.106). Again, FMT had the largest effect on viral composition with lesser but significant effects of FFT relative to CON irrespective of administration route (Supplementary table S2). Interestingly, as opposed to bacteria, the Shannon diversity index of mucosa-associated viruses increased following either FMT or FFTo but not FFTr relative to CON (both p < 0.05, Fig. 3B). The same pattern was observed for viral Shannon diversity index in the luminal compartment. Opposite to what was observed for bacteria, co correlation between mucosal viral Shannon diversity index and small intestinal NEC severity was observed (data not shown). The majority of identifiable viruses across the groups were prokaryotic viruses (phages) primarily belonging to the order Caudovirales (e.g. Siphoviridae, Myoviridae, Podoviridae; Supplementary figure S5), although several eukaryotic viruses were also identified but in low relative abundance. Remarkably, while FMT led to mucosal and luminal enrichment of eukaryotic virus families (e.g. Retroviridae, Herpesviridae) relative to CON, this was not the case for FFT treatment.

### Host gut mucosal transcriptome profiling

To gain insight into the host treatment response, we then performed RNA-Seq on standardized ileal mucosa samples - biological replicates of the samples used for mucosal microbiota analysis. Initially, a plotting of principal components showed that the FMT group separated from the remaining groups along the second principal component (Fig. 4A). Notably, for the two FFT groups we found no single differentially expressed gene relative to CON (FDR adjusted p < 0.10), whereas FMT increased the expression of 86 genes and decreased the expression of 41 genes compared with CON (Fig. 4B-C). When applying fold-change criteria (log2 > 1), 29 and 16 known genes were up- and downregulated by FMT relative to CON, respectively (Supplementary table S3). A network analysis of differentially expressed genes identified *TLR4* and *CD14*, as well as *HCK*, a gene involved in neutrophil degranulation as key genes in the FMT-enriched network, whereas ING2, a gene involved in apoptosis, as well as interferon-induced genes such as IFIT1 and OASL were among the key genes in the diminished network (Fig. 4D). Functional annotation of differentially expressed genes in FMT vs. CON mucosa showed that the most significantly affected pathways were related to immune activation and host defense mechanisms. Interestingly, FMT upregulated genes involved in bacterial response pathways and downregulated genes related with viral response (Supplementary table S4).

**Figure 4.**
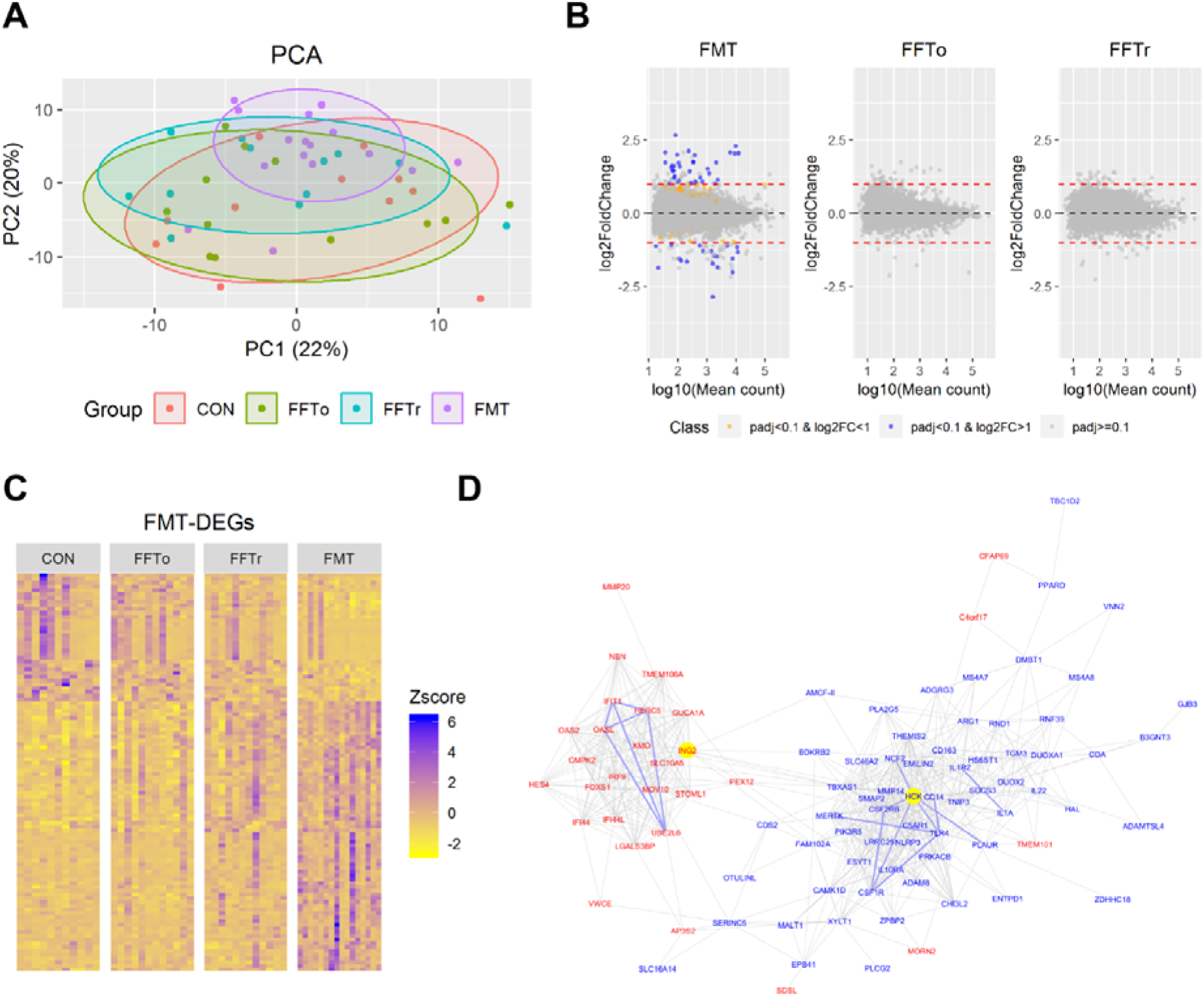
Host gut mucosal transcriptome profiling following FMT or FFT treatment. A. Principal component analysis plot of global gene expression differences based on RNA Seq. data. B. Volcano plots of pairwise comparisons with the CON group, highlighting differentially expressed genes based on false discovery rate-adjusted statistics and fold-change criteria. C. Z-score based gene expression heat map showing differentially expressed genes of the FMT vs. CON pairwise comparison. D. Gene network cluster showing FMT-upregulated genes in blue and FMT-downregulated genes in red.

### Systemic immune cell characterization

Finally, we measured the levels of basic immune cell subtypes in the bloodstream both shortly after intervention and at euthanasia to investigate any induction of systemic immunity. On day 3, briefly after the final treatment administration, we found marginally increased neutrophil counts in all intervention groups relative to CON (all p < 0.05, Supplementary figure S6), whereas monocyte and total lymphocyte levels were not affected. However, the helper T cell (CD4^+^CD8^-^) and naïve T cell fraction (CD4^-^CD8^-^) were increased and decreased, respectively in FMT relative to FFTr (both p < 0.05). Two days later, these differences had all disappeared.

## DISCUSSION

Despite being recognized for decades, NEC remains a clinical challenge today. Currently, disease prophylaxis is limited to the use of breastfeeding as well as probiotics, which lack standard recommendations and is subject to scrutiny^37^. The treatment of suspected NEC consists of enteral feeding discontinuation, enteral or parenteral antibiotics and symptomatic medical treatment. Still, almost 10% of extremely preterm infants develop NEC, and among these, the risk of death or disability is high^38^. Moreover, the widespread use of antibiotics in preterm infants due to suspected infection is related with an increase in antibiotics resistance^39^, while animal experiments indicate that neonatal antibiotics perturb immune development in a microbiota-dependent manner and increase the risk of secondary infections^40,41^. Collectively, there is a need for more effective therapeutic options with less collateral impact.

In this experiment, we aimed to test the NEC-preventive effect of bacteria-free FFT with an additional focus on preclinical safety. Using cesarean-delivered preterm pigs, the most ambitious and clinically relevant animal model of NEC, we showed 1) a proof of principle for FFT treatment with a slight advantage of oral vs. rectal administration route 2) an inconspicuous FFT safety profile, whereas cognate FMT treatment was associated with a range of adverse effects.

This study is the first to describe the use of FFT as a means to protect against NEC, and among the first to describe the therapeutic potential of FFT altogether. A small case report described the successful use of FFT in recurrent *Clostridioides difficile-infected* patients, the only patient group where FMT is routinely used. Besides ensuring clinical remission, FFT changed the gut bacterial and viral composition within each patient^18^. Recently, FFT from lean mice donors was shown to change the bacterial and viral gut microbiota, in turn reducing weight gain and improving glucose tolerance in diet-induced obese mice^19^. Besides, FFT improves the reshaping of antibiotics-induced microbiota disruption^42^. These studies all administered FFT in the proximal gut (oral, gastric, jejunal), while we additionally assessed the effects of gastric vs. rectal administration, and although only modestly different, the effect of gastric administration was superior in reducing pathological severity. Only Rasmussen *et al* studied isolated virus particles after column adsorption^19^, whereas the remaining studies including the current one studied sterile fecal filtrate containing viruses as well as secretome and metabolome, which contain many bioactive substances. Hence, to confirm causality of the donor fecal virome, follow-up experiments investigating isolated virus fractions and inactivated viruses are needed to unequivocally show the effect of the virus fraction.

Whereas no adverse effects were observed during and following FFT treatment, we identified several safety issues related with FMT treatment. This study was intended to demonstrate non inferiority of FFT compared with FMT in terms of NEC development, since we previously found a robust reduction in NEC incidence following FMT, using the same batch of donor material and comparable study design^15^. In the previous FMT experiment, combined gastric and rectal administration increased sepsis-related mortality, whereas exclusively rectal administration appeared safe. We later confirmed that exclusively oral FMT administration caused 90% of animals to become lethally septic and with inner organs colonized by FMT-derived *Escherichia coli* (unpublished results). In the current experiment, rectal FMT, which we perceived as effective and safe, failed to prevent NEC, reduced total body growth and was detrimental to the small intestine by reducing structural integrity and function capacity. This suggests the existence of a severity spectrum where the presence of viable bacteria and in turn gastric administration increases the risk of adverse effects. However, upon bacterial removal, oral administration becomes safe, as shown in the present study.

Bacteriophage density is increased in the mucus layer relative to the lumen^43^, which inspired us to investigate effects of treatment on the mucosa-associated microbiota. Whereas all groups contained Enterobacteriaceae as part of the luminal microbiota, we found that CON and to some extent FMT animals had an increased abundance of this bacterial family in their mucosa, which has consistently been linked with NEC development in preterm infants^11,13,14,44^. Moreover, the diversity of mucosa-associated bacteria correlated with NEC severity. Meanwhile, oro-gastric FFT increased mucosa-associated viral diversity. Hence, a plausible mechanism of action for FFT might be a particular enrichment of phages in the mucus layer, which in turn reduce the density of bacteria e.g. Enterobacteriaceae in close proximity to the mucosa. In the case of FMT, administering a large dose of viable bacteria concomitantly, might dilute or abolish this effect.

A peculiar finding concerned the colonization of recipients with *Streptococcus* and *Lactobacillus*, which together constituted a major fraction of the donor microbiota. While these genera were absent in the gut of control animals, *Streptococcus* was the second most abundant in FFT animals, whereas *Lactobacillus* was hardly detectable. Contrarily, *Lactobacillus* was the second most abundant genus in FMT animals, a confirmation of previous observations^15^, while *Streptococcus* relative abundance was reduced. Whether this dichotomy is of any clinical relevance, remains to be seen. Regardless, it might be due to competitive exclusion between these two closely related genera, and could reflect the phage specificity in the FFT solution that either inhibit *Lactobacillus* or favor *Streptococcus* growth.

A major concern and potential obstacle for the use of FFT in preterm infants is the risk of transferring eukaryotic viruses capable of infecting human cells from an older donor individual into a compromised recipient. In this study, phages, particularly those belonging to the order Caudovirales, were dominating across groups and anatomical niches. The second most abundant virus could not be identified, but as the five other most abundant viruses (totaling 98% abundance) were all phages, and since phage database coverage is much lower than for eukaryotic viruses, likely it is an uncharacterized phage. Indeed, Caudovirales is the dominating phage in infants as well^16,17^, and as importantly, very few eukaryotic viruses inhabit the new born as well as the one month old infant gut, whereas the abundance increases at four months^17^. Regardless, we found that FMT but not FFT increased the abundance of a number of clinically relevant mammalian viruses e.g. Herpesviridae. Either the sterile filtering process inhibits a fraction of the largest virus particles from entering the FFT solution, or *in vivo* conditions following FFT prevent these viruses from infecting host cells. In any case, this aspect is important for FFT clinical feasibility.

To increase the understanding of how gut microbiota directed therapies may act or interact to prevent intestinal pathology, we performed a global gene expression analysis of the host gut mucosa. This analysis was performed in anatomically standardized tissue specimens, which were largely devoid of severe pathology. Hence, the results should be interpreted as isolated treatment effects with minimal bias due to pathological state. Strikingly, no gene expression levels were significantly affected by FFT relative to CON, which indicates that no particular interaction occurs between administered viruses (or small molecules) and host immune cells, implying that FFT exerts its effect on bacteria only. Direct interaction between phages and host via TLR9 signaling has been demonstrated under germ-free conditions^45^, but under physiological conditions, this effect may be of negligible importance. On the other hand, FMT treatment regulated mucosal gene expression levels towards increased immune response to bacteria and concomitant reduction in host response to viruses. Furthermore, FMT led to a transient induction of circulating helper T cells shortly after administration. From a safety perspective, the lack of collateral effects on host immunity favors the use of FFT over FMT.

As the FFT concept has been demonstrated in the porcine species only, and using just a single batch of donor fecal material, a generalized conclusion across species and individual donors in terms of safety and efficacy cannot be made yet. Regardless, in light of these findings and the unique potential, we encourage FFT safety and feasibility testing in preterm infants using a rigid safety paradigm. Donor and stool should be subject to an extensive screening procedure, equivalent to existing pediatric FMT procedures^46^, and bacteria removed using validated techniques^20^. Furthermore, the data indicates the use of a prophylactic approach with early neonatal treatment, and favors the use of oro-gastric administration route. Besides, we advise against the use of FMT in preterm infants.

