## Supplementary material for "Fecal filtrate transfer protects against necrotizing enterocolitis in preterm pigs"

**SUPPLEMENTAL MATERIAL**

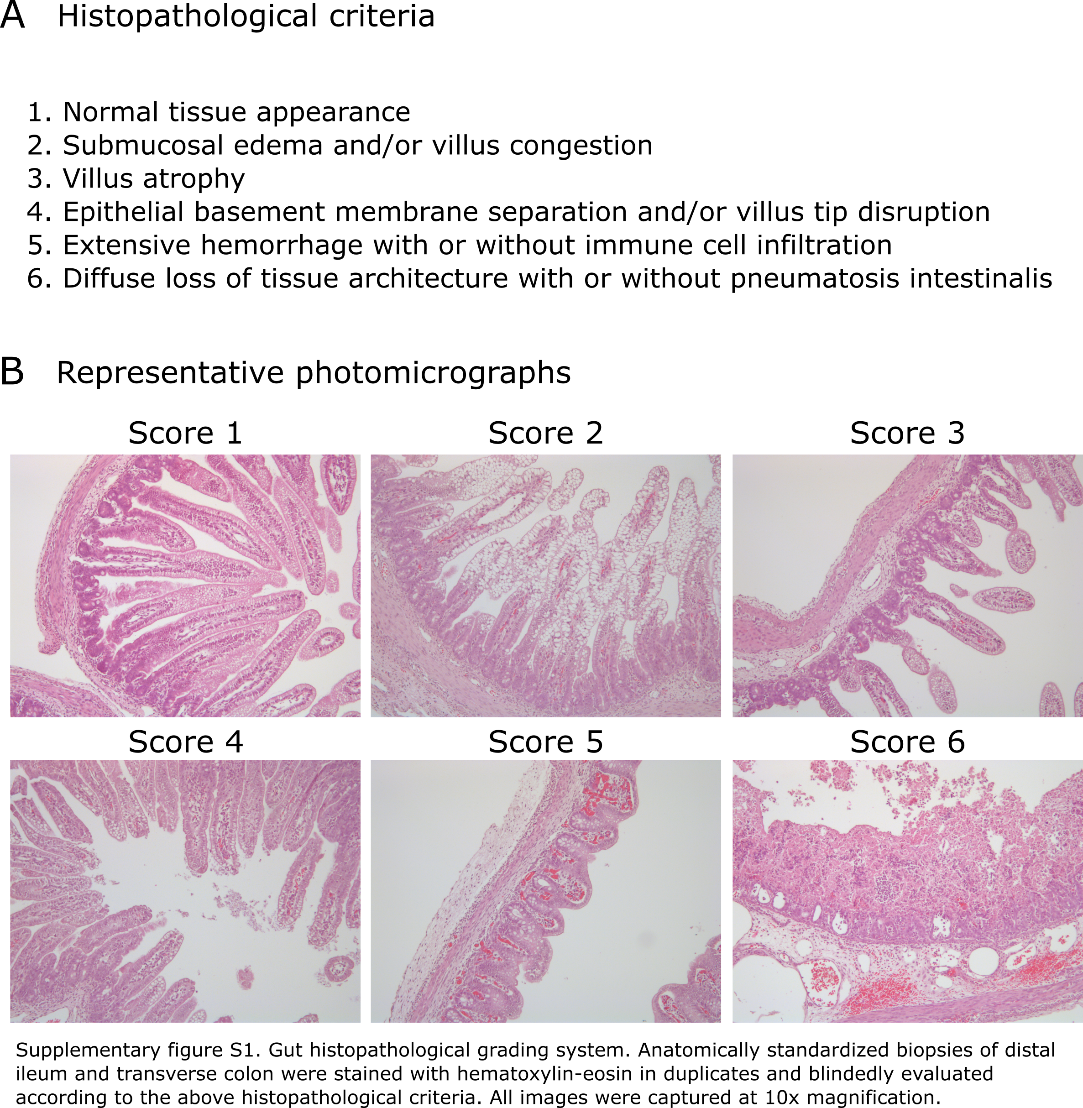

**Supplementary figure S1. Presentation of gut histopathological grading system**. A. Definition of the six grades of increasing histopathological severity. B. Representative micrographs illustrating each severity grade.

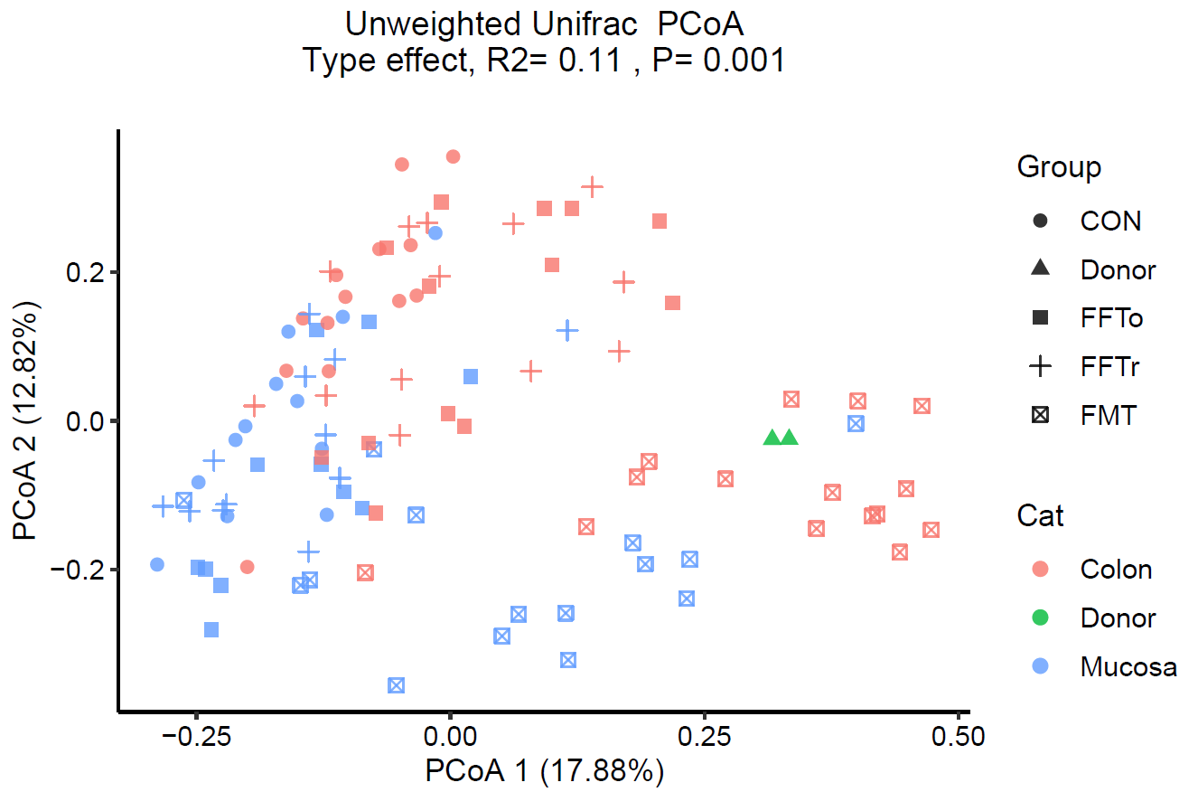

**Supplementary figure S2. Bacterial composition in different gut niches**. Comparison of bacterial composition in the gut mucosal and luminal niche presented as principal component analysis plot based on unweighted UniFrac dissimilarity.

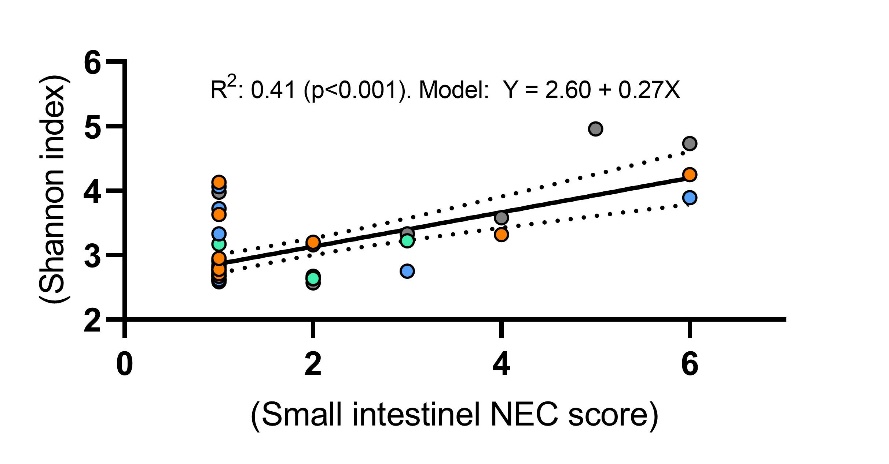

**Supplementary figure S3. Association between macroscopic NEC severity of the small intestine and mucosa bacterial diversity as assessed by Shannon index**.

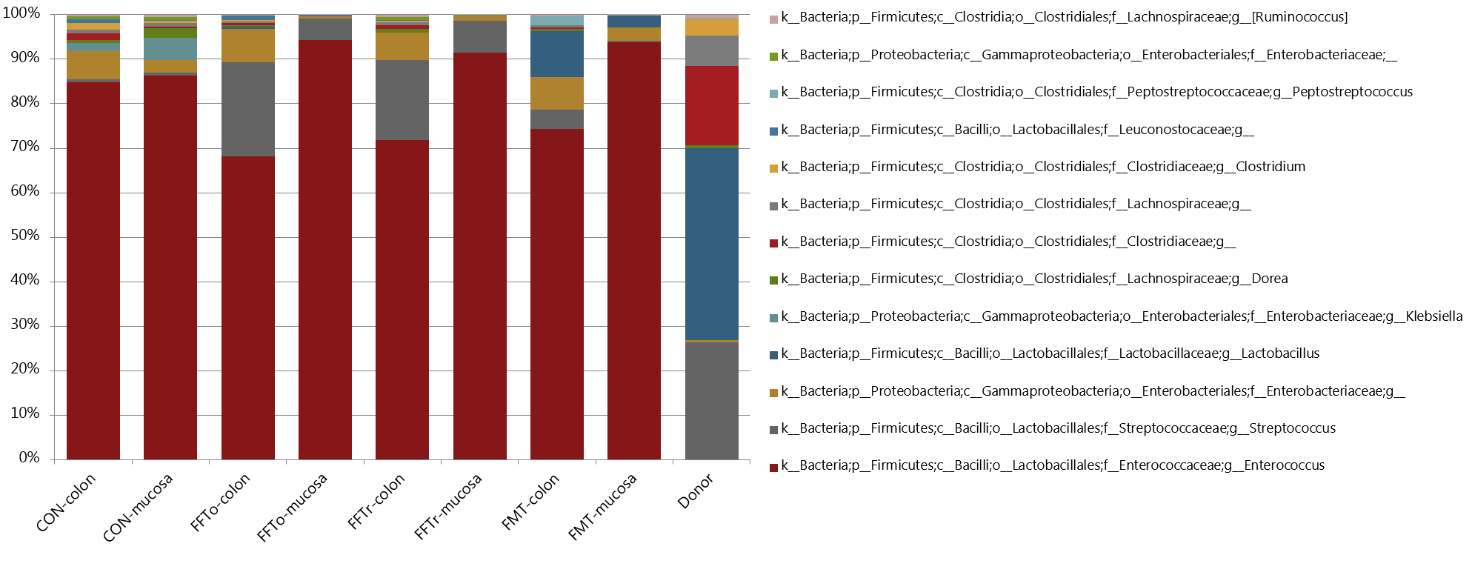

**Supplementary figure S4. Bacterial taxonomy of gut mucosa and lumen**. Relative abundance of bacteria based on 16S rRNA gene amplicon sequencing and summarized at genus level. Genera with more than 1% relative abundance across groups are included. Bacterial composition of donor fecal material is included as reference.

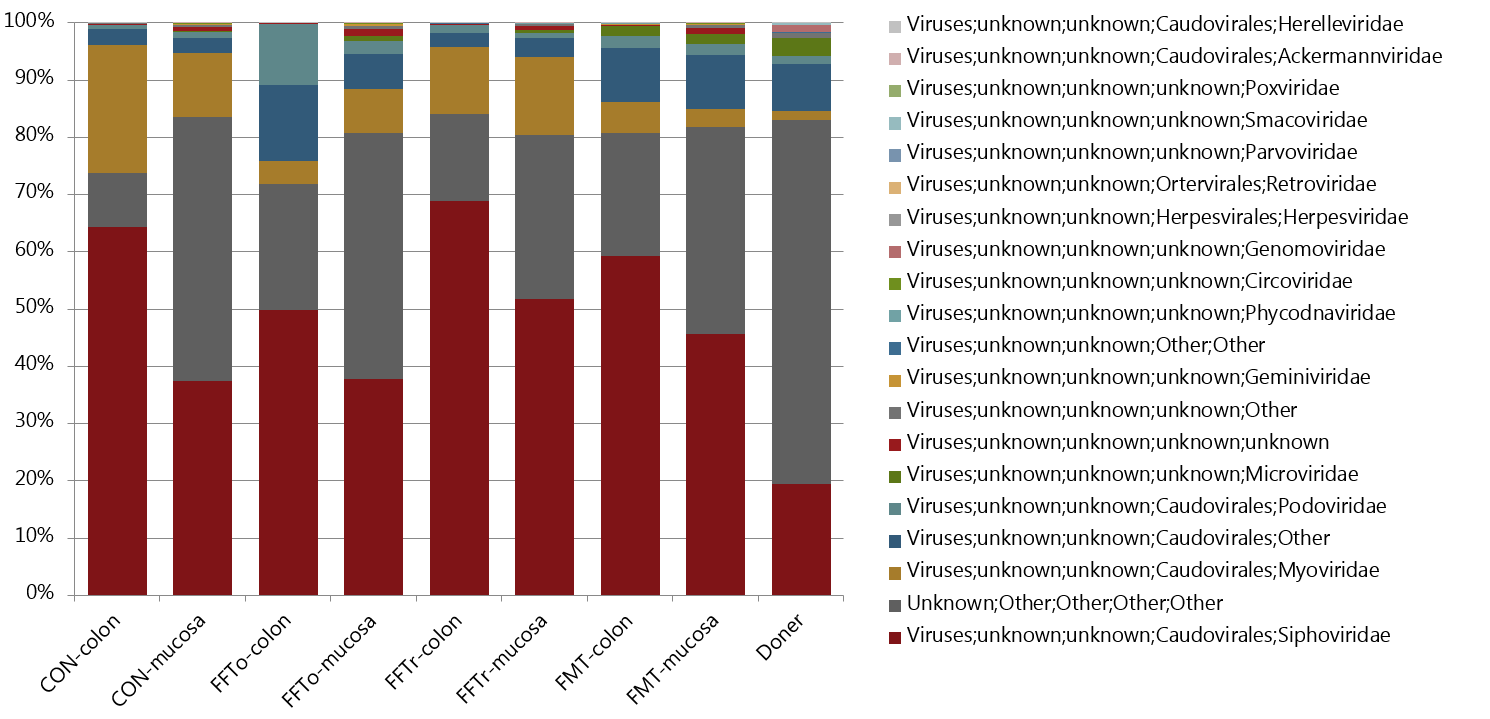
 **Supplementary figure S5. Viral taxonomy of gut mucosa and lumen**. Relative abundance of viruses based on metagenome sequencing of virus-enriched and bacteria-depleted fecal samples, summarized at family level. Viral composition of donor fecal material is included as reference.

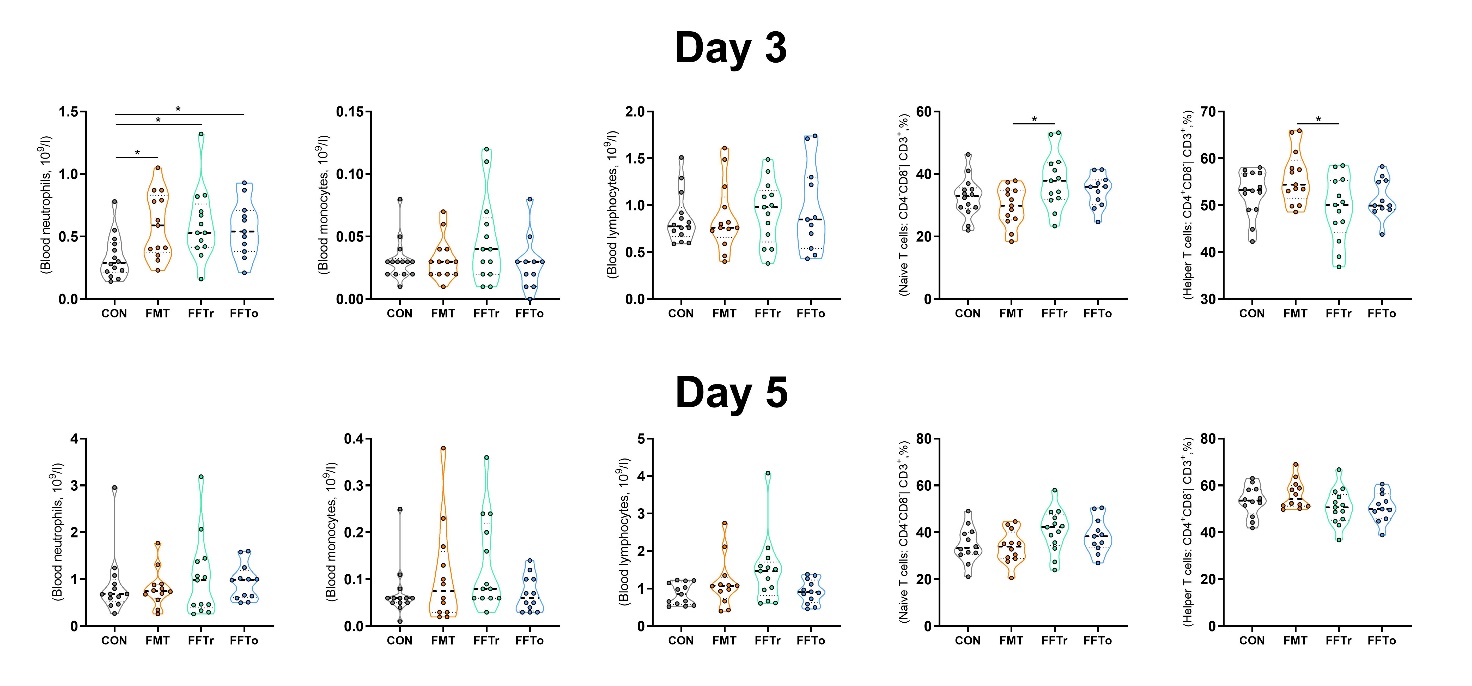

**Supplementary figure S6. Systemic immune cell characterization following FMT or FFT treatment.** Blood neutrophils, monocytes and total lymphocytes as well as helper T cell and naïve T cell fractions were measured on day 3 shortly after treatments and again on day 5. * denotes a statistical probability level below 0.05.

**Supplementary table S1.** Infant formula composition

| **Constituent** | **Infant formula** |
| --- | --- |
| Fantomalt | 40 g/L |
| Seravit | 12 g/L |
| Miprodan 40 | 15 g/L |
| Lacprodan DI-9224 | 60 g/L |
| Liquigen | 75 g/L |
| Calogen | 20 g/L |

**Supplementary table S2**. Pairwise comparisons of beta diversity.

|  | CON vs FMT | CON vs FFTr | CON vs FFTo | FMT vs FFTr | FMT vs FFTo | FFTr vs FFTo |
| --- | --- | --- | --- | --- | --- | --- |
| Mucosa associated bacteria | | |  |  |  |  |
|  | 6.66 | 2.03 | 3.52 | 6.61 | 5.54 | 1.41 |
|  | (0.002) | (0.022) | (0.002) | 0.002 | 0.002 | 0.131 |
| Luminal bacteria | | |  |  |  |  |
|  | 14.0 | 3.69 | 4.83 | 10.2 | 8.60 | 1.30 |
|  | (0.001) | (0.001) | (0.001) | (0.001) | (0.001) | (0.187) |
| Mucosa associated viruses | | |  |  |  |  |
|  | 1.66 | 0.73 | 1.23 | 1.47 | 1.63 | 1.28 |
|  | (0.09) | (0.65) | (0.27) | (0.13) | (0.08) | (0.22) |
| Luminal viruses | | |  |  |  |  |
|  | 6.17 | 2.67 | 4.06 | 1.64 | 2.41 | 1.72 |
|  | (0.002) | (0.017) | (0.005) | (0.094) | (0.011) | (0.066) |
| PERMANOVA test for pairwise differences in beta diversity. Bacterial diversity is measured in unweighted UniFrac dissimilarity and viral diversity using Bray-Curtis dissimilarity. F-values with corresponding FDR adjusted q-value in parenthesis. | | | | | | |

**Supplementary table S3**. Mucosal RNA seq. differentially expressed genes.

| **Gene** | **CON** | **FMT** | **FFTr** | **FFTo** | **log2FoldChange (FMT/CON)** | **FDR adj. P-value** |
| --- | --- | --- | --- | --- | --- | --- |
| ARG1 | 27 | 181 | 67 | 17 | 2,66 | 0,05 |
| TGM3 | 3859 | 18896 | 8240 | 4811 | 2,29 | 0,02 |
| MMP9 | 99 | 426 | 218 | 122 | 2,12 | 0,04 |
| MT1A | 17 | 83 | 17 | 12 | 2,07 | 0,08 |
| IL22 | 37 | 166 | 77 | 70 | 2,00 | 0,05 |
| DUOXA1 | 150 | 665 | 367 | 217 | 1,98 | 0,06 |
| CDA | 158 | 544 | 398 | 184 | 1,75 | 0,04 |
| MMP13 | 192 | 607 | 210 | 227 | 1,75 | 0,05 |
| PLA2G5 | 18 | 64 | 32 | 21 | 1,74 | 0,02 |
| DUOX2 | 1111 | 3695 | 2009 | 937 | 1,73 | 0,07 |
| MS4A7 | 58 | 197 | 61 | 73 | 1,73 | 0,05 |
| VNN2 | 38 | 117 | 45 | 34 | 1,63 | 0,00 |
| GJB3 | 35 | 112 | 77 | 50 | 1,62 | 0,05 |
| IL1R2 | 64 | 192 | 100 | 72 | 1,60 | 0,03 |
| HAL | 39 | 108 | 41 | 33 | 1,55 | 0,07 |
| CD163 | 611 | 1836 | 759 | 671 | 1,53 | 0,04 |
| IL1A | 37 | 106 | 53 | 49 | 1,45 | 0,04 |
| CHI3L2 | 42 | 116 | 64 | 52 | 1,45 | 0,04 |
| RBP4 | 860 | 2018 | 819 | 810 | 1,36 | 0,07 |
| TNIP3 | 51 | 124 | 79 | 48 | 1,26 | 0,04 |
| AMCF-II | 361 | 872 | 556 | 439 | 1,23 | 0,04 |
| ADGRG3 | 49 | 114 | 50 | 61 | 1,21 | 0,03 |
| SOCS3 | 1585 | 3582 | 2134 | 1648 | 1,15 | 0,04 |
| RNF39 | 76 | 172 | 85 | 63 | 1,12 | 0,07 |
| MS4A8 | 467 | 1036 | 593 | 567 | 1,09 | 0,04 |
| PRSS22 | 281 | 598 | 363 | 319 | 1,07 | 0,07 |
| SLC46A2 | 56 | 117 | 114 | 87 | 1,06 | 0,01 |
| RND1 | 192 | 402 | 251 | 246 | 1,03 | 0,08 |
| CSF2RB | 1282 | 2679 | 1817 | 1445 | 1,00 | 0,03 |
| C4orf17 | 88 | 40 | 66 | 57 | -1,09 | 0,09 |
| HES4 | 972 | 368 | 387 | 599 | -1,11 | 0,08 |
| UBE2L6 | 6569 | 2951 | 4838 | 6180 | -1,12 | 0,07 |
| IRF9 | 25 | 12 | 15 | 21 | -1,14 | 0,04 |
| FOXS1 | 1992 | 775 | 1541 | 1680 | -1,30 | 0,05 |
| SDSL | 1197 | 482 | 427 | 951 | -1,30 | 0,04 |
| LGALS3BP | 16240 | 6317 | 10578 | 15171 | -1,34 | 0,09 |
| CMPK2 | 2555 | 901 | 1636 | 2330 | -1,46 | 0,09 |
| IFI44 | 3375 | 1159 | 2032 | 2903 | -1,48 | 0,10 |
| GUCA1A | 151 | 58 | 98 | 147 | -1,56 | 0,08 |
| OAS2 | 11636 | 3977 | 6700 | 9993 | -1,56 | 0,07 |
| MMP20 | 332 | 137 | 162 | 225 | -1,66 | 0,05 |
| HERC5 | 3820 | 1037 | 1895 | 3079 | -1,85 | 0,10 |
| VWCE | 40 | 8 | 11 | 32 | -1,87 | 0,03 |
| IFIT1 | 13239 | 3346 | 6123 | 9729 | -2,02 | 0,07 |
| OASL | 2565 | 611 | 1331 | 2280 | -2,86 | 0,08 |
| Differentially expressed genes in CON vs FMT with at log2 fold change > 1 and FDR adjusted test probability level < 0.10 summarized in descending order according to fold change level. | | | | | | |

**Supplementary table S4**. RNA seq. pathway enrichment analysis.

| **Term ID** | **Term description** | **Gene count** | **Background** | **FDR adj. P-value** |
| --- | --- | --- | --- | --- |
| GO:0051707 | response to other organism | 25 | 835 | 3,99E-09 |
| GO:0006952 | defense response | 26 | 1234 | 6,30E-07 |
| GO:0002237 | response to molecule of bacterial origin | 14 | 317 | 9,75E-07 |
| GO:0031347 | regulation of defense response | 19 | 676 | 1,27E-06 |
| GO:0009617 | response to bacterium | 17 | 555 | 2,39E-06 |
| GO:0032496 | response to lipopolysaccharide | 13 | 298 | 2,76E-06 |
| GO:0009605 | response to external stimulus | 30 | 1857 | 4,93E-06 |
| GO:0051704 | multi-organism process | 33 | 2222 | 5,14E-06 |
| GO:0009615 | response to virus | 12 | 270 | 6,40E-06 |
| GO:0002252 | immune effector process | 20 | 927 | 1,61E-05 |
| GO:0006950 | response to stress | 40 | 3267 | 1,61E-05 |
| GO:0006954 | inflammatory response | 14 | 482 | 5,43E-05 |
| GO:0006955 | immune response | 25 | 1560 | 7,33E-05 |
| GO:0051607 | defense response to virus | 9 | 181 | 1,10E-04 |
| GO:0002682 | regulation of immune system process | 23 | 1391 | 1,20E-04 |
| GO:0002376 | immune system process | 31 | 2370 | 1,40E-04 |
| GO:0080134 | regulation of response to stress | 22 | 1299 | 1,40E-04 |
| GO:0071222 | cellular response to lipopolysaccharide | 8 | 146 | 1,80E-04 |
| GO:0032101 | regulation of response to external stimulus | 16 | 732 | 2,00E-04 |
| GO:0033993 | response to lipid | 17 | 825 | 2,00E-04 |
| Top 20 gene ontology terms in CON vs FMT summarized according to FDR adjusted test probability level. | | | | |
